# Mapping phenotypic heterogeneity in melanoma onto the epithelial-hybrid-mesenchymal axis

**DOI:** 10.1101/2022.04.05.485702

**Authors:** Maalavika Pillai, Gouri Rajaram, Pradipti Thakur, Nilay Agarwal, Srinath Muralidharan, Ankita Ray, Jason A Somarelli, Mohit Kumar Jolly

## Abstract

Epithelial to mesenchymal transition (EMT) is a well-studied hallmark of epithelial-like cancers that is characterized by loss of epithelial markers and gain of mesenchymal markers. Interestingly, melanoma, which is derived from melanocytes of the skin, also undergo phenotypic plasticity toward mesenchymal-like phenotypes under the influence of various micro-environmental cues. Our study connects EMT to the phenomenon of de-differentiation (i.e., transition from proliferative to more invasive phenotypes) observed in melanoma cells during drug treatment. By analyzing 78 publicly available transcriptomic melanoma datasets, we found that de-differentiation in melanoma is accompanied by upregulation of mesenchymal genes, but not necessarily a concomitant loss of an epithelial program, suggesting a more “one-dimensional” EMT that leads to a hybrid epithelial/ mesenchymal phenotype. Samples lying in the hybrid epithelial/mesenchymal phenotype also correspond to the intermediate phenotypes in melanoma along the proliferative-invasive axis - neural crest and transitory ones. Interestingly, as melanoma cells progress along the invasive axis, the mesenchymal signature does not increase monotonically. Instead, we observe a peak in mesenchymal scores followed by a decline, as cells further de-differentiate. This biphasic response recapitulates the dynamics of melanocyte development, suggesting close interactions among genes controlling differentiation and mesenchymal programs in melanocytes. Similar trends were noted for metabolic changes often associated with EMT in carcinomas in which progression along mesenchymal axis correlates with the downregulation of oxidative phosphorylation, while largely maintaining glycolytic capacity. Overall, these results provide an explanation for how EMT and dedifferentiation axes overlap with respect to their transcriptional and metabolic programs in melanoma.

## Introduction

Epithelial to mesenchymal transition (EMT) is a well-characterized phenomenon involved in multiple axes of cancer progression, such as metastasis and drug resistance. EMT is commonly associated with morphological changes, functional changes (increased migration, invasion, and immune invasion (Chakraborty et al., 2021; Hanahan & Weinberg, 2011; Sahoo et al., 2021) and molecular changes, including upregulation of EMT markers and transcription factors (TFs), such as *VIM, ZEB1, SNAI1* and *TWIST1*. While the phenomenon of EMT has largely been characterized for epithelial cancers (such as breast cancer and lung adenocarcinoma), similar molecular, functional and morphological changes have also been observed in non-epithelial cancers, such as sarcomas (Somarelli, Shetler et al. 2016, Jolly, Ware et al. 2019), glioblastoma (Siebzehnrubl et al., 2013), myeloma (Roccaro et al., 2015), lymphoma (Lemma et al., 2013; Sánchez-Tilló et al., 2013), leukemia (Stavropoulou et al., 2016; Wu et al., 2018) and melanoma (Kahlert et al., 2017) in preclinical and clinical settings.

Standard-of-care chemotherapy for melanoma often includes targeted inhibition of BRAF or MEK signaling. While these targeted agents provide clinical benefit to melanoma patients, resistance to these therapies is common. Therapy-resistant melanomas often undergo de-differentiation, which is characterized by loss of melanocytic markers such as *MLANA, TRPM1* and *TYR* and gain of invasive molecular markers such as *c-JUN, NGFR* and *ZEB1* (Denecker et al., 2014; Fallahi-Sichani et al., 2017; Rambow et al., 2018; Ramsdale et al., 2015). The de-differentiation trajectory of melanoma cells is characterized by a transition along the proliferation-invasion axis, from a melanocytic phenotype to an undifferentiated phenotype while passing through the intermediate transitory and neural crest stem cell-like (NCSC) phenotypes (**Fig. 1A**). This trajectory is the reverse of the differentiation that occurs during melanocyte development, where undifferentiated tissue in the embryonic neural plate give rise to highly migratory and mesenchymal neural crest cells, some of which differentiate into melanocytes upon reaching the epidermis (Dupin & le Douarin, 2003). Therapy resistant melanoma is also commonly associated with a mesenchymal-like phenotype with more invasive and aggressive features (Denecker et al., 2014; Ramsdale et al., 2015; Su et al., 2017; Vandamme et al., 2020; Wouters et al., 2020). These relationships between de-differentiation, invasion, and EMT pathways in response to therapy suggest EMT and de-differentiation programs in melanoma may be linked.

**Fig 1.**
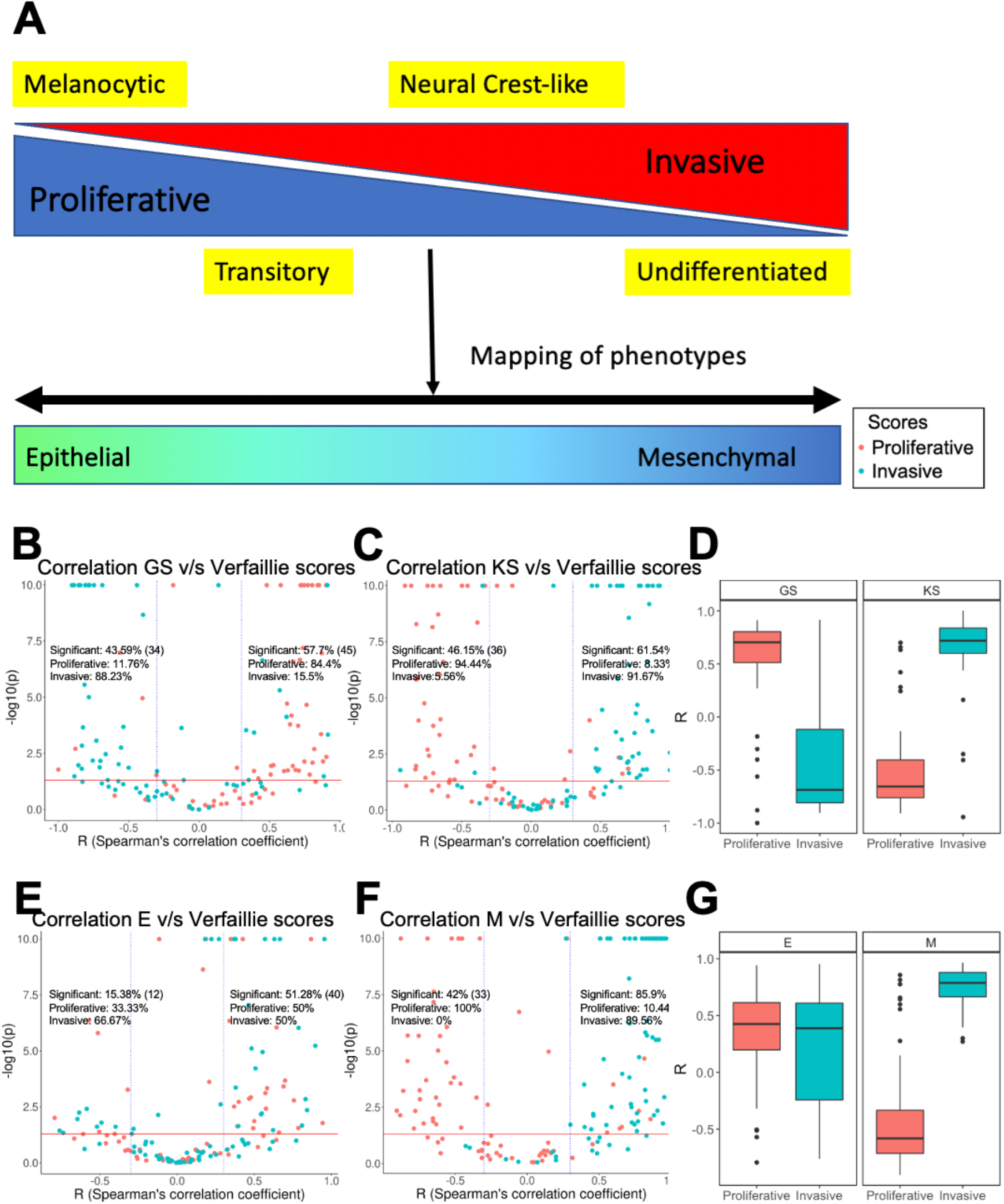
Mapping phenotypic heterogeneity in melanoma onto the EMT axis. **A**. A schematic representation. Volcano plots depicting Spearman’s correlation coefficient and -log_10_(p-value) of 78 datasets for Verfaillie proliferative and invasive gene set with **B**. 76GS EMT scoring metric, and with **C**. KS EMT scoring metric **D**. Boxplot depicting range of correlation coefficients for KS and 76GS with Verfaillie invasive and proliferative gene sets. Volcano plots depicting the Spearman’s correlation coefficient and -log_10_(p-value) of 78 datasets for Verfaillie proliferative and invasive gene set with **E**. Epithelial gene set (E scores) and **F**. Mesenchymal gene set (M scores). **G**. Boxplot depicting range of correlation coefficients for E and M scores with Verfaillie invasive and proliferative gene sets. Inset labelled “Significant” is calculated as the fraction of datasets (out of 78) which show a significant correlation trend (r < - 0.3 or r > 0.3, p < 0.05). Absolute number of such significant points (datasets) for the specified cut of is mentioned in brackets. “Proliferative” and “Invasive” labels represent the percentage of significant correlations that are between the EMT score and proliferative score or invasive score, respectively.

The similarity between EMT and de-differentiation programs extends beyond cell-intrinsic alterations and impacts cell-extrinsic changes as well. EMT often leads to varied extracellular matrix (ECM) stiffness and density (Deng et al., 2021; Fattet et al., 2020; Kumar et al., 2014) and altered cell-matrix and cell-cell interactions (Bianchi et al., 2010; Kilinc et al., 2021). In melanoma, acquisition of de-differentiated and invasive phenotypes is often accompanied with changes in composition and physical properties of ECM, and modified cell-matrix interactions and cell morphology (Kaur et al., 2019; Long et al., 2019; Spoerri et al., 2021). Increased expression of matrix metalloproteases (MMPs), immune evasion (characterized by both signaling-mediated immune suppression (e.g. by TGF-ß release) and prevention of immune cell entry into tumors by dense collagen matrix/low α-SMA expression), increased inflammatory markers (such as TNF-α, NF-kB and AP-1) and cytoskeleton remodeling have been closely linked to the acquisition of an invasive phenotype and loss of melanocytic differentiation regulator MITF (Dilshat et al., 2021; Jensen et al., 2018; M. H. Kim et al., 2016; Lal et al., 2013; Miskolczi et al., 2018; Riesenberg et al., 2015). All of these changes are reported with EMT progression as well in multiple epithelial cancers (Radisky & Radisky, 2010; Suarez-Carmona et al., 2017; Tripathi et al., 2016). Such extensive similarity between EMT and de-differentiation programs in cancer-microenvironment cross-talk and niche construction underscore the potential of common regulatory pathways involved in both EMT and de-differentiation.

Another common feature that links EMT in epithelial cancers to de-differentiation in melanoma is the presence of intermediate or hybrid phenotypes. Hybrid epithelial/mesenchymal (E/M) cells express molecular and functional characteristics of both epithelial (high proliferation and cell-cell adhesion, low invasion) and mesenchymal (low proliferation and cell-cell adhesion, high invasion) cells (Jolly et al., 2022). On the other hand, melanoma intermediate phenotypes, which comprise transitory and neural crest-like stem cell-like (NCSC) phenotypes, exhibit combined features of proliferative and invasive phenotypes (Hoek et al., 2008; Tsoi et al., 2018) (**Fig. 1A**). Gene regulatory networks for EMT and melanoma provide a mechanistic basis for explaining the existence of these hybrid/intermediate states (Jolly et al., 2016; Pillai & Jolly, 2021). An overlap in key regulators and stabilizers for hybrid E/M phenotypes and melanoma phenotypes (such as *ZEB1, NFATC2, CDH1, SNAI2, NRF2*) suggest common regulatory links (Bocci et al., 2019; Caramel et al., 2013; Denecker et al., 2014; Gupta et al., 2005; Jessen et al., 2020; Perotti et al., 2019; Subbalakshmi et al., 2020, 2022). For instance, *SNAI2*, a stabilizer of the hybrid E/M phenotype, is a key regulator of the NCSC phenotype and metastasis in melanoma, suggesting its involvement in regulating the intermediate phenotypes in melanoma as well (Gupta et al., 2005; Subbalakshmi et al., 2022). However, intriguingly, certain regulators show opposite trends in melanoma and EMT. For instance, *ZEB2* is considered an inducer of EMT in epithelial cancers, but in the context of melanoma, it inhibits the mesenchymal phenotype (Vandamme et al., 2020; Vandewalle et al., 2005). Other molecules that show opposite effects include *KLF4* (Subbalakshmi et al., 2021; Zhang et al., 2018) and *TFAP2A* (Campbell et al., 2021; Dimitrova et al., 2017). Thus, understanding the mechanistic underpinning of how the de-differentiation and EMT programs are linked can help decipher reasons for the similarities and differences between these pathways across cancers.

In this study, we map the de-differentiation axis in melanoma (also called proliferative-invasive/ P-I axis) to the EMT axis using previously defined scoring metrics (Byers et al., 2013; Sahoo et al., 2021; Subramanian et al., 2005; Tan et al., 2014). We compare the extent to which a gain in a mesenchymal signature corresponds to a loss in the epithelial signature during de-differentiation of melanoma. By deciphering the interdependencies between de-differentiation and mesenchymal programs, the differences in molecular regulation between EMT and de-differentiation can be explained. We have identified that the mesenchymal program, but not the epithelial program, is closely linked to de-differentiation. Although the mesenchymal signature enrichment shows a strong negative correlation with a differentiated/melanocytic transcriptional program, it does not increase monotonically during de-differentiation. This non-monotonic trend is also captured by metabolic programs associated with EMT, such as glycolysis and HIF1α, but not with metabolic programs associated with differentiation/melanocytic genes, such as the MITF-regulated OXPHOS pathway. Our results indicate that phenotypic heterogeneity in melanoma occurs along a proliferative-invasive axis that correlates with a “one-dimensional EMT” in which cells transition along a mesenchymal axis without an alteration in epithelial phenotype. Moreover, our analyses support a conceptual model in which mesenchymal scores initially increase but then saturates/ decreases as melanoma cells undergo dedifferentiation, thus suggesting caution with respect to timescales considered in *in vitro* analysis of drug-induced dedifferentiation programs. Deciphering such inter-connections among multiple axes of plasticity in a cancer cell population may guide potent combinatorial therapeutic strategies aimed at controlling transitions to a more hybrid cell type with combined features of both proliferation and invasion.

## Materials and Methods

### Software and Datasets

Publicly available datasets from Gene Expression Omnibus (GEO), The Cancer Genome Atlas (TCGA), Cancer Cell Line Encyclopedia (CCLE-Broad Institute) (Barretina et al., 2012), and National Cancer Institute-60 (NCI-60) databases were analyzed. Microarray data were downloaded from GEO using GEOquery Bioconductor R package. All analyses done on R version 4.1.0. *ggplot2*, and *ggpubr* R packages were used to create and customize plots.

### Pre-processing of Datasets

Microarray datasets, with un-mapped probe IDs, were pre-processed by mapping the probe IDs onto their gene symbols using the relevant platform annotation table. In the case of multiple probes mapping to the same gene, the mean expression of all the probes was considered for that gene. For non-normalized RNA-Seq datasets TPM normalization followed by *log2* transformation with an offset value of 1 was used.

### ssGSEA

Single-sample Gene Set Enrichment Analysis, an extension of Gene Set Enrichment Analysis (GSEA) (Barbie et al., 2009; Subramanian et al., 2005), calculates separate enrichment scores for each sample and a gene set. Each score represents the degree to which genes in a gene set are up or down-regulated in a sample. We calculated ssGSEA scores for the Verfaillie proliferative and Verfaillie invasive gene sets (Verfaillie et al., 2015), Hoek proliferative and Hoek invasive gene sets (Hoek et al., 2008), the epithelial (E) and mesenchymal (M) gene sets of the EM tumor gene signature genes and cell lines gene signatures in the KS scoring metric (Tan et al., 2014), and the Tsoi melanocytic, transitory, NCSC, and undifferentiated gene set (Tsoi et al., 2018).

### Calculation of EMT Scores

We calculated EMT scores of datasets using four metrics-76 Gene Signature (76GS), Kolmogorov -Smirnov test (KS), E score and M score. 76GS and KS were calculated as defined earlier (Byers et al., 2013; Chakraborty et al., 2020; Tan et al., 2014). 76GS score is a metric for how epithelial a sample is, i.e., higher scores reflect greater association with an epithelial phenotype. The KS score is a metric for how mesenchymal a sample is. The higher the KS score of a sample, the greater is its association with a mesenchymal phenotype. While 76GS scores do not have a predefined range of scores, KS scores lie within a +1 to -1 range. The E and M scores are ssGSEA scores for epithelial and mesenchymal gene lists, respectively, for the KS scoring metric (Sahoo et al., 2021). For calculating KS, E and M scores, datasets were classified based on whether the samples were derived from cell-lines or tumors and the appropriate gene sets were used.

### Correlations

All correlation values were calculated using Spearman’s correlation coefficient, unless mentioned otherwise. Spearman’s correlation coefficient method generates a coefficient ranging between –1 to +1, where +1 indicates a strong positive correlation, and –1 indicates a strong negative correlation between two variables. It determines the correlation between any monotonically related variables-linear or non-linear. Correlations with R >0.36 and p<0.05 are considered significant.

### Moving Window Average

A moving window average is used to quantify the gradient for a variable along a given axis. A window covering 60% of the entire range of the axis is created and the average value of the variable for all samples in the window is calculated. Then the window is then shifted by 1% and the average is re-calculated. This is iteratively repeated to cover the entire range. The slope of the averages determines the magnitude and direction of the gradient.

### Conditional probabilities

Once the cell lines were sorted into their respective phenotypes and the conditional probabilities were obtained, the statistical significance and p-values for the conditional probabilities were calculated using the one-proportion Z test.

The z-score was calculated using the equation

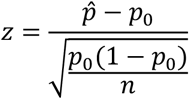

where 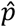 is the observed proportion, *p*_0_ is the null probability, and *n* is the sample size. The obtained value of z was then converted into the corresponding p-value using the standard normal distribution. If the obtained p-value < 0.05, it was considered significant.

### Assigning phenotypes to samples

In order to identify samples belonging to the 4 phenotypes (melanocytic, transitory, NCSC and undifferentiated), we calculated ssGSEA scores based on gene sets for each of these phenotypes (Tsoi et al., 2018). Samples lying in the top 10% scores were assigned that particular phenotype.

### Metabolic scores

The oxidative phosphorylation (OXPHOS) and glycolysis (Glyco) scores in our study were calculated using ssGSEA carried out with the corresponding hallmark gene sets for these pathways (obtained from Molecular Signature Database (MSigDB) (Liberzon et al., 2015)). The HIF-1 signature - which is a surrogate for glycolysis - was quantified based on a method previously reported (Yu et al., 2017). This method uses expression levels of their downstream target genes to capture the respective enzyme activities. A total of 84 downstream genes for AMPK and 59 downstream genes for HIF-1 were used and the scores were obtained using the Singscore method performed on these gene sets (Foroutan et al., 2018; Muralidharan et al., 2022). The fatty acid oxidation (FAO) scores were calculated based on the equation previously reported (Jia et al., 2020) which uses expression levels of 14 FAO enzyme genes.

## Results

### Enrichment of mesenchymal genes can capture the extent of de-differentiation in melanoma

To test whether EMT and de-differentiation in melanoma programs are correlated with one another, we used previously-defined EMT scores – KS and 76GS (Byers et al., 2013; Tan et al., 2014) – and ssGSEA scores for Verfaillie proliferative and invasive (Verfaillie et al., 2015) and Hoek proliferative and invasive (Hoek et al., 2008) melanoma gene sets and investigated their correlation coefficients across 78 datasets. Also, to dissect the contributions of epithelial and mesenchymal gene set separately, we calculated the ssGSEA scores (Barbie et al., 2009; Subramanian et al., 2005) for corresponding gene sets individually too (Tan *et al*. 2014), referred here as E and M scores respectively (Sahoo et al., 2021). A sample with a higher 76GS or E score is more epithelial while a higher KS or M score refers to more mesenchymal phenotype. Thus, given the overlap between mesenchymal and invasive programs, we expected invasive scores to correlate positively with KS and M scores and negatively with 76GS and E scores. We also expected opposite trends for proliferation scores: negative correlations with KS and M scores and positive correlation with 76GS and E scores. We visualized the relationships between these pathways as volcano plots in which each dot corresponds to a dataset analysed. For positively correlated metrics, we expect the majority of data sets to lie in the top right rectangle, while those displaying a significant negative correlation are expected to lie in the top left rectangle.

34 out of 78 datasets (43.59%) showed a significant negative correlation (r < - 0.3, p < 0.05) between 76GS and one of the two Verfaillie (proliferative, invasive) scores. In 30 out of those 34 datasets (88.23%), 76GS scores correlated negatively with invasive scores, while in remaining 4 datasets (11.76%), 76GS scores correlated negatively with proliferative scores (**Fig. 1B**, left). Similarly, among 45 datasets that showed a positive correlation (r > 0.3, p < 0.05) between 76GS scores and one of Verfaillie scores, 38 (84.4%) cases had a positive correlation between 76GS and proliferative scores, and in the remaining seven datasets, 76GS scores correlated positively with invasive scores (**Fig. 1B**, right). Overall, both the scoring metrics (76GS and KS) displayed clear correlation with Verfaillie and Hoek proliferative and invasive scores across 78 datasets in the expected direction, with KS showing fewer false positive cases (cases where the correlation is significant but in a direction opposite to the expected one) as compared to 76GS (**Fig. 1B-D, S1A-C**).

Because gain of mesenchymal features is reported more commonly in melanoma as compared to loss of epithelial features, we decoupled the epithelial and mesenchymal components of the scoring metrics (E and M scores, respectively). The KS method provides information on genes that are associated with an epithelial phenotype and those with a mesenchymal state separately. Using the genes from the KS scoring method we segregated the genes and calculated individual ssGSEA scores for epithelial and mesenchymal gene lists and re-evaluated their correlation with proliferative and invasive scores in melanoma. While epithelial genes continued to show random distributions of samples throughout the plot, mesenchymal genes showed clear segregation of proliferative and invasive scores based on Spearman’s correlation coefficients, i.e., invasive scores were positively correlated with M score while proliferative scores were negatively correlated with the M scores (**Fig. 1E-G, S1D-F**). This observation suggests that mesenchymal genes, but not epithelial genes, can capture the phenotypic heterogeneity displayed by melanoma along the proliferative-invasive axis.

For further analysis, we focused only the Verfaillie gene sets, because it has high levels of overlap with gene sets for the intermediate phenotypes that were previously identified (Tsoi et al., 2018) (**Fig. S1G**). Thus, a continuous scoring metric defined for the Verfaillie gene set is expected to be more sensitive for capturing intermediate phenotypes as compared to the Hoek gene set.

Because correlation coefficients only provide an overall trend in data, we wished to determine how proliferative and invasive scores vary along the E and the M axis. For this purpose, we generated two dimensional EMT plots of the data sets in which E and M scores are represented along each of the two axes. These plots display how epithelial or mesenchymal a given sample is (Barbie et al., 2009; Sahoo et al., 2021; Subramanian et al., 2005). We then overlay information on the proliferative and invasive scores for each sample. As expected, across various datasets, proliferative and invasive scores for samples had a stronger visible gradient along the M axis as compared to the E axis (**Fig. 2A-B**). To quantify this gradient, we used a rolling window to estimate the increase of average proliferative and invasive scores across the E and M axis. For this, we start with a rolling window covering 60% of the entire range along a given axis and calculate the average proliferative (P) or invasive (I) score within that window. Then the window is shifted by 1% and the average is re-calculated. This process is repeated until the entire range is covered, and the change in averages is plotted. For an axis that strongly correlates with the change in scores, we expect a steeper slope. The nature of a slope (positive or negative) is determined by the correlation between the axis and the score. Both axes trend in the expected direction, with a positive slope for invasive scores and negative slope for proliferative scores along the M axis and vice versa for the E axis (**Fig. 2C**). This analysis also reveals that the M axis has a steeper curve than the E axis for both P and I scores. These results suggest that proliferative-invasive heterogeneity in melanoma can be considered as a “one-dimensional form” of EMT where the mesenchymal program enrichment increases as cells become more invasive, but the epithelial program need not be suppressed concomitantly, as often tacitly assumed for the case of EMT.

**Fig 2.**
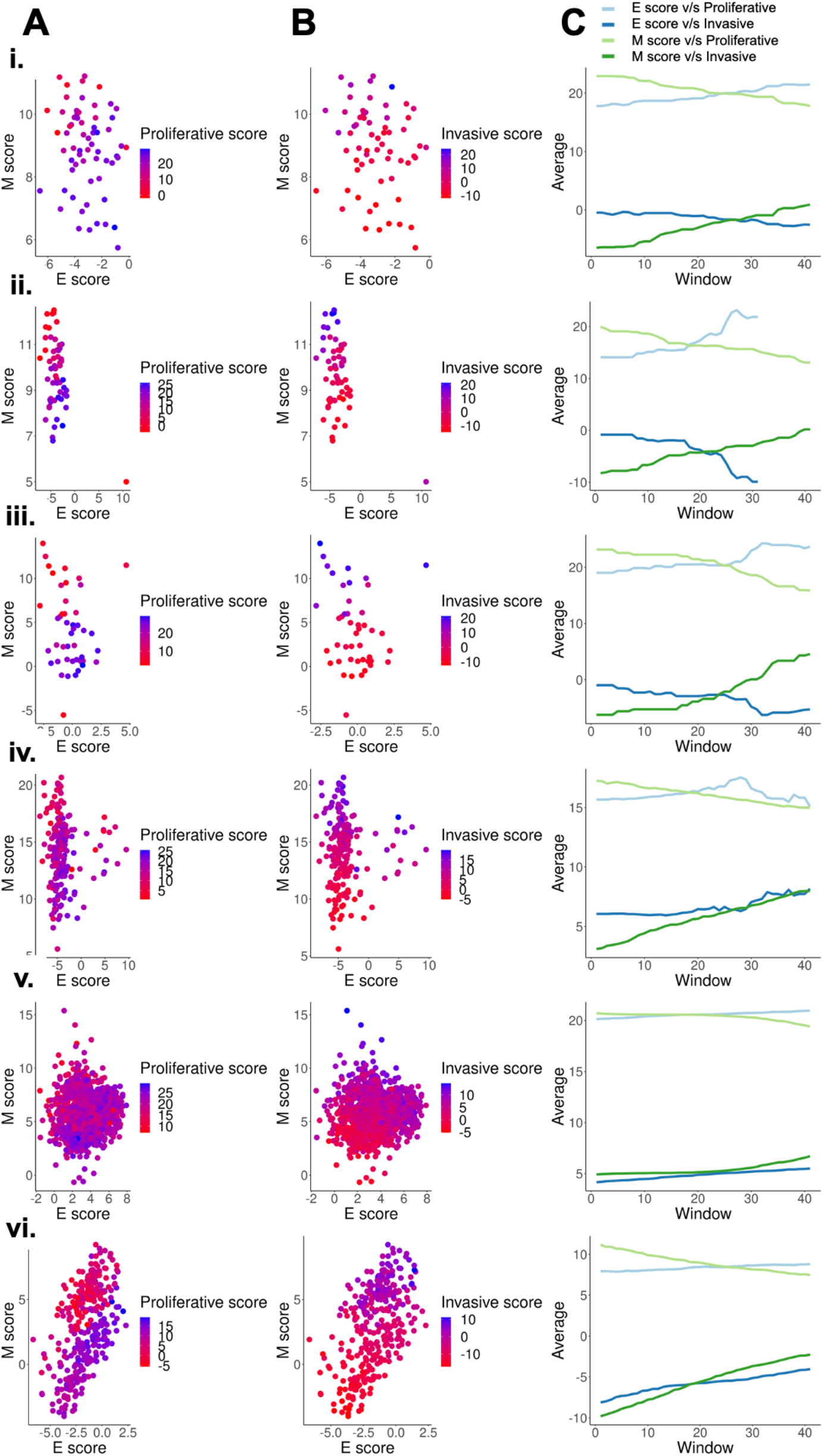
Scoring metrics based on mesenchymal genes capture de-differentiation better than metrics based on epithelial genes. Two dimensional EMT plots of different types of datasets-i. GSE7127 (63 melanoma cell lines - microarray), ii. CCLE (59 cell lines - microarray), iii.GSE4843 (45 tumor samples - microarray), iv.GSE65904 (214 tumor samples - microarray),v. GSE72056 (1257 single-cell tumor samples), vi.GSE81383 (307 single-cell tumor sample) depicting the variation of **A**. Proliferative scores along the E and M score axes. **B**. Invasive scores along the E and M score axes. **C**. Quantifying the proliferative and invasive score gradient along the E-M axes using a rolling window.

### The mesenchymal axis follows a non-monotonic relationship with de-differentiation

Because the M score axis was able to capture the phenomenon of de-differentiation quantified by continuous scoring metrics, such as the proliferative and invasive scores, we next tested if the discretized phenotypes also arrange themselves in order of appearance along the two dimensional EMT plane. The classification of samples into four categories - melanocytic, transitory, neural crest-like and undifferentiated (in order of increasing de-differentiation) - for GSE80829, GSE10916, GSE4843, GSE7127 and GSE116237 was done as previously defined (Pillai & Jolly, 2021; Rambow et al., 2018). Along the proliferative-invasive plane, samples displayed a strong negative relation between the two scores, i.e., proliferative scores of samples decreased as their invasive score increased. The four phenotypes also appeared in the expected order (Su et al., 2017; Tsoi et al., 2018), with the melanocytic samples having the highest proliferative scores and lowest invasive scores, and the undifferentiated samples displaying the lowest invasive scores and highest proliferative scores (**Fig. 3A**). However, the two dimensional EMT plane failed to resolve the four phenotypes in terms of these four phenotypes showing non-overlapping scores. Since the E score axis performed poorly previously (**Fig. 1E,G**) in these metrics, we quantified the ability of M score axis alone to resolve the four phenotypes by quantifying the conditional probability of a sample to belong to the intermediate phenotypes (transitory and NCSC), given that they lie in an intermediate M score range. Interestingly, samples with intermediate M scores were significantly likely to belong to the transitory phenotype (**Fig. 3B, S3A, Table 1**). However, the probability of these samples to belong to the NCSC phenotype was negligible. In some datasets (GSE7127, GSE116237), the melanocytic phenotype was also significantly enriched in the intermediate M score populations. However, the melanocytic phenotype cells were enriched in the bottom M score population as well, and were not uniquely present in the intermediate score range like the transitory phenotype cells (**Fig. S3B-D**).

**Table 1.** Conditional probabilities for a sample belonging to a particular phenotype given that it lies in the intermediate M score range.

| Dataset | P(Melanocytic\|<br>Intermediate M<br>Score) | p-value | P(Transitory\|<br>Intermediate M<br>Score) | p-value | P(NCSC\| Intermediate<br>M Score) | p-value | P(Undifferentiated\|<br>Intermediate M<br>Score) | p-value |
| --- | --- | --- | --- | --- | --- | --- | --- | --- |
| GSE80829 | 0.17 | 0.8 | 0.43 | 0.02 | 0 | 1 | 0.39 | 0.06 |
| GSE7127 | 0.48 | 0.01 | 0.43 | 0.02 | 0 | 1 | 0.09 | 0.96 |
| GSE11623<br>7 | 0.36 | 0 | 0.49 | 0 | 0.09 | 1 | 0.06 | 1 |

**Fig 3.**
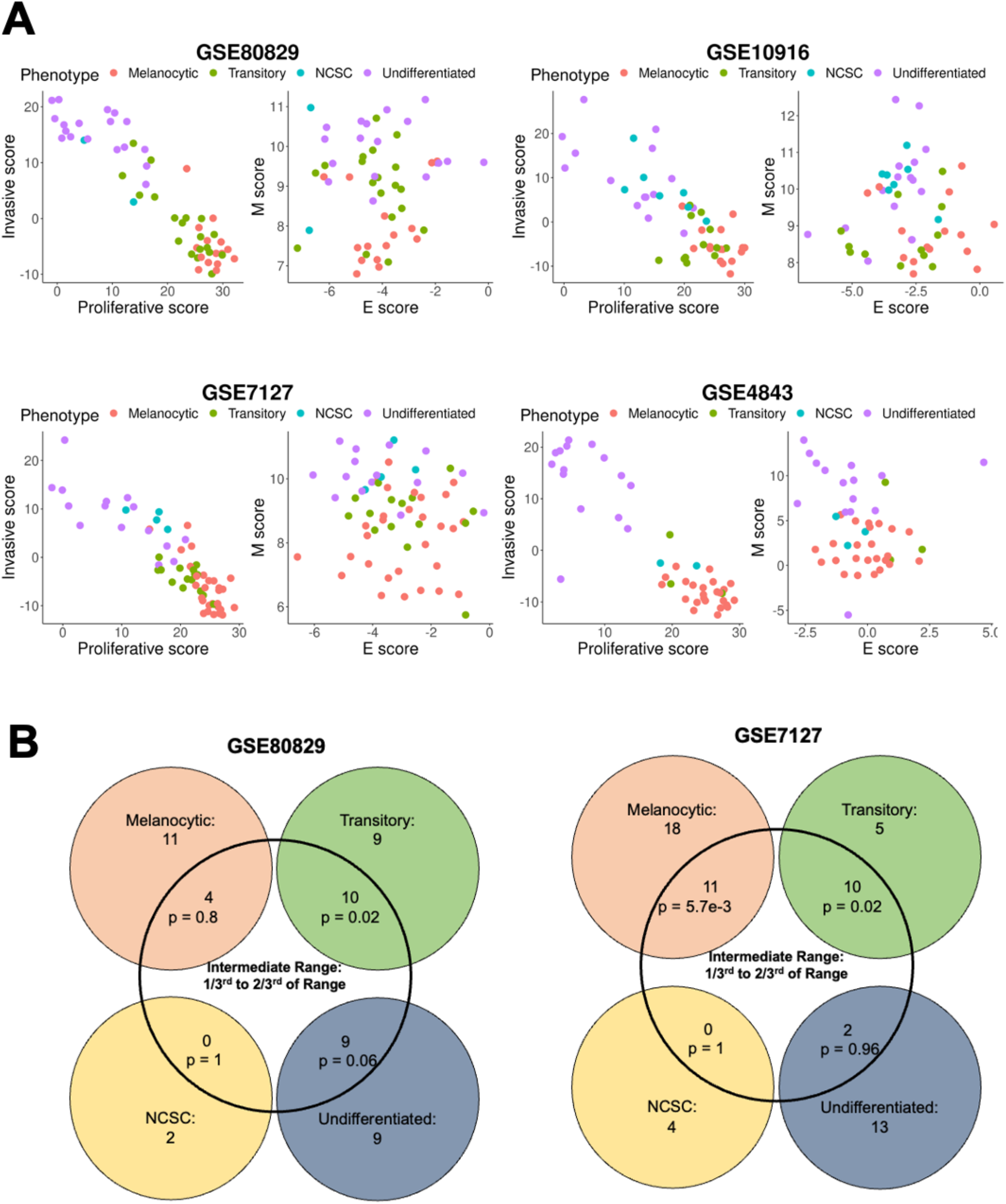
Variation of the four molecular phenotype scores along the epithelial, mesenchymal, proliferative, and invasive axes. **A**. Plotting samples classified into four phenotypes onto the E-M, proliferative-invasive score axes. **B**. Venn diagram depicting the intersection of the four phenotype scores of samples and intermediate M scores. p represents p-value for the conditional probability that a sample belongs to the phenotype given that they lie in the intermediate M score range.

To further dissect the relationship between the four phenotypes and the M score axis, we quantified the change in M score with respect to the invasive scores for the four phenotypes. To identify the four phenotypes, we used ssGSEA scores for gene sets defined for each of the four phenotypes (Tsoi et al., 2018). The top 10% of samples that had the highest scores for a particular gene set, were assigned the label of that particular phenotype. Interestingly, we observed that in these samples there was a non-monotonic increase in M scores as invasive score/de-differentiation increased. As samples progressed from NCSC to undifferentiated, M scores either decreased (**Fig. 4C-E**) or remained the same (**Fig. 4A-B, 4F**). In the context of melanocyte development, neural crest cells are precursors for melanocytes with high migratory potential and high levels of EMT markers (Dupin & le Douarin, 2003; Tang et al., 2020; Wessely et al., 2021). Thus, the nonmonotonic increase in the mesenchymal program seen here is reminiscent of the differentiation of melanocytes.

**Fig 4.**
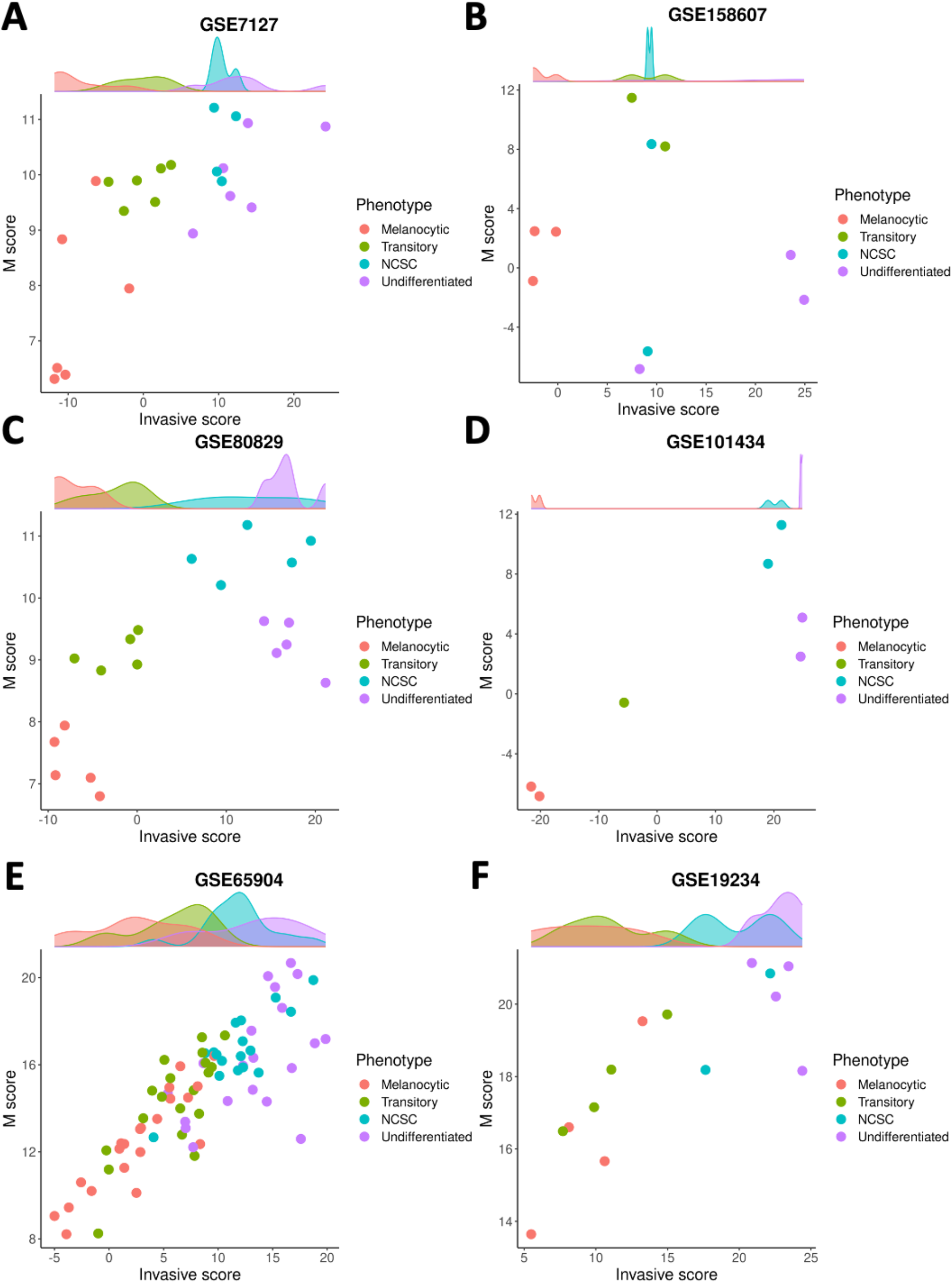
The mesenchymal axis follows a non-monotonic relationship with de-differentiation. Plotting M scores against invasive scores for different phenotypes along the P-I axis in many datasets: **A**. GSE7127 **B**. GSE158607 **C**. GSE80829 **D**. GSE101434 **E**. GSE65904 **F**. GSE19234

### Metabolic reprogramming along the proliferative-invasive axis in melanoma

The EMT status of epithelial cancer cells is often associated with distinct metabolic programs. Generally speaking, EMT is negatively correlated with the enrichment of oxidative phosphorylation (OXPHOS) and fatty acid oxidation (FAO), but positively correlated with glycolysis (Muralidharan et al., 2022). In melanoma, the proliferative state is associated with high levels of OXPHOS and the invasive phenotype is associated with high levels of glycolysis (Abildgaard & Guldberg, 2015; Bettum et al., 2015; Gelato et al., 2017; Laurenzana et al., 2017), reinforcing the commonalities between these two different instances of phenotypic plasticity. Computational analysis has suggested the existence of four metabolic sub-populations (Jia et al., 2020): 1) OXPHOS-high/ glycolysis-low, 2) OXPHOS-low/ glycolysis-high, 3) OXPHOS-low/glycolysis-low, and 4) OXPHOS high/glycolysis-high.

To assess whether the OXPHOS-glycolysis metabolism axis can be mapped onto the proliferation-invasion axis, we calculated Spearman’s correlation coefficients between the metabolic scores (OXPHOS and glycolysis) and the de-differentiation scores (proliferative and invasive scores) (**Fig 5A-C**) across the 78 datasets. In 36 out of 78 datasets where the OXPHOS scores correlate significantly with proliferative scores, 32 datasets show a positive correlation. Similarly, among 43 datasets showing a significant correlation of OXPHOS scores with invasive scores, all of them showed negative correlation. Thus, overall, OXHOS scores corelated positively with proliferative scores and negatively with invasive scores (**Fig 5A**). Glycolysis scores, on the other hand, did not show a clear relationship with EMT status, with a subset of datasets showing trends in both the directions (positive and negative correlation) both for proliferative and invasive scores (**Fig 5B**). This difference is reminiscent of prior observations for the association of EMT with OXPHOS and glycolysis in which glycolysis is only moderately correlated with EMT status, but OXPHOS is consistently negatively correlated with EMT (Muralidharan et al., 2022). This trend is substantiated by observations that in cases where OXPHOS is positively correlated with proliferative scores or negatively correlated with invasive scores, glycolysis scores do not show any particular direction of enrichment with either proliferative or invasive axes (**Fig 5C**).

**Fig 5.**
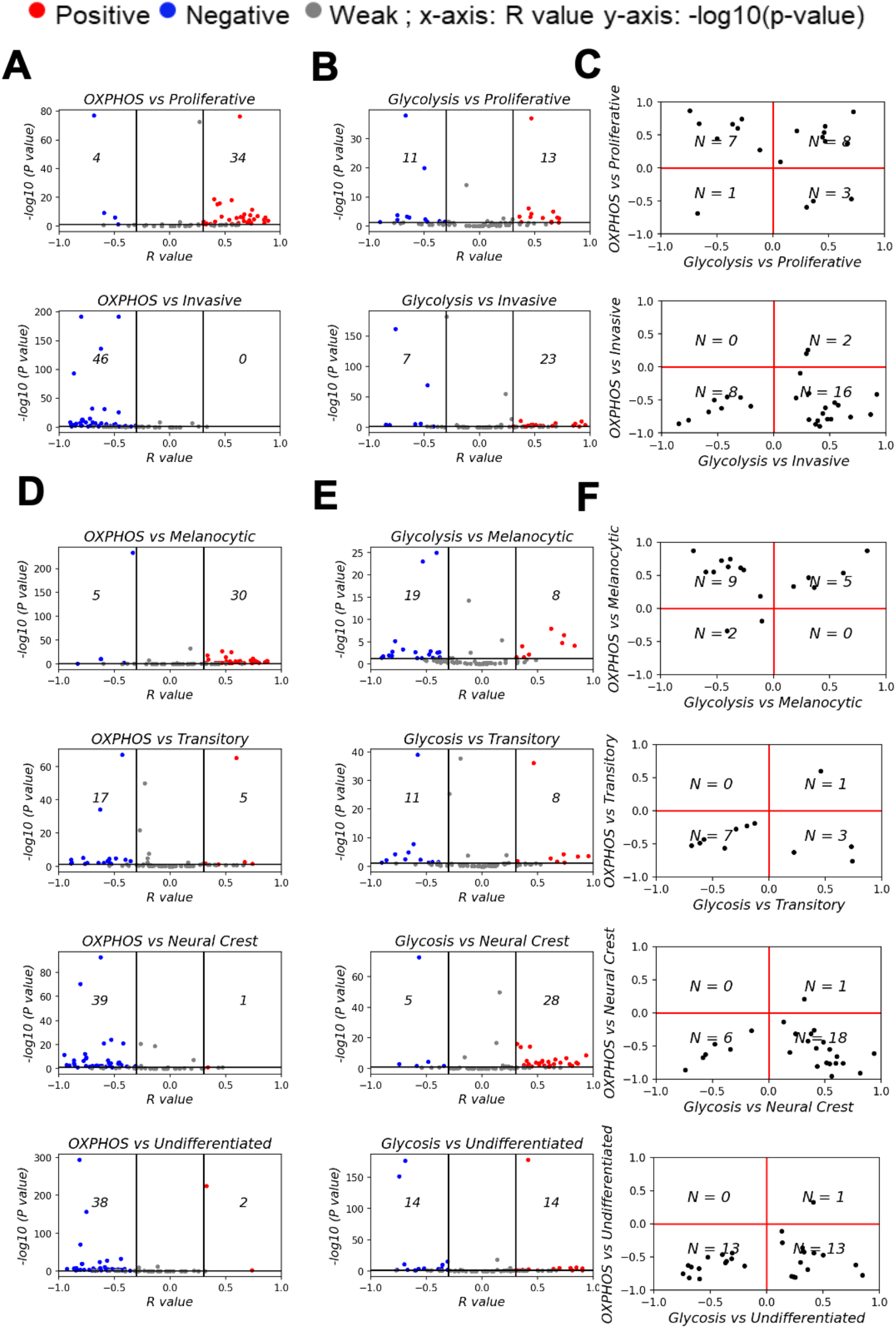
Mapping metabolic programs associated with EMT onto the de-differentiation program axes. Volcano plots depicting Spearman’s correlation coefficient and -log_10_(p-value) of 78 datasets for **A**. Hallmark OXPHOS and Verfaillie gene set. **B**. Hallmark glycolysis and Verfaillie gene set. **C**. Spearman’s correlation coefficient between OXPHOS and Glycolysis and Verfaillie scores. **D**. Hallmark OXPHOS and Tsoi gene set. **E**. Hallmark glycolysis and Tsoi gene set. **F**. Spearman’s correlation coefficient between OXPHOS and Glycolysis and Tsoi scores. N represents number of samples present in a given quadrant

We next sought to dissect whether intermediate melanoma phenotypes might be enriched for a specific metabolic profile. To investigate this trend, we calculated the Spearman’s correlation coefficients for metabolic scores and ssGSEA scores for gene signatures corresponding to each of the four molecular phenotypes of melanoma (**Fig 5D-F**). OXPHOS showed a clear shift from datasets with a significant positive correlation with a melanocytic phenotype to a significant negative correlation for the undifferentiated phenotype (**Fig 5D**). On the contrary, glycolysis scores do not show a clear shift from negative to positive correlations with de-differentiation (**Fig 5E**). Similar to the non-monotonic trend observed for M-scores, the glycolysis scores show the strongest positive correlation trends for the NCSC phenotype. Undifferentiated phenotype scores have a mixture of positively correlated and negatively correlated datasets with respect to glycolysis scores. Put together, these observations suggest that the regulatory modules controlling the switch to glycolysis are likely linked to the mesenchymal program rather than the de-differentiation one. On the other hand, regulatory modules for OXPHOS are likely to be closely linked to the melanocytic differentiation program. This trend is in accordance with experimental evidence that suggests that OXPHOS in melanoma cells is regulated by PGC1α, a downstream target of MITF, a key regulator of melanocyte differentiation (Haq et al., 2013; Vazquez et al., 2013). Interestingly, fatty acid oxidation, which is also directly controlled by MITF via SCD (Vivas-García et al., 2020), also displays trends similar to OXPHOS (**Fig. S4A**) while a HIF1α signature, that is commonly associated with the invasive phenotype follows a non-linear trend similar to glycolysis (**Fig. S4B**), suggesting that it is linked to the mesenchymal program rather than the de-differentiation program.

## Discussion

De-differentiation in melanoma occurs in response to targeted therapy. This process may be mediated by transitions across a spectrum of phenotypes in which melanocytic cells treated with BRAF/MEK inhibitors pass through a transitory phenotype, followed by the NCSC phenotype, before becoming completely un-differentiated (Rambow et al., 2018; Su et al., 2017; Tsoi et al., 2018). This trajectory is accompanied by loss of a proliferative signature and gain of invasive characteristics. Here, we decipher the relationship between de-differentiation and EMT in melanoma. These processes are often considered to co-occur during drug treatment (Fallahi-Sichani et al., 2017; Ramsdale et al., 2015; Riesenberg et al., 2015); however, comparison of EMT and de-differentiation scores reveal that the two processes may be more closely related to the mesenchymal program rather than the loss of an epithelial-like state or an EMT program *per se*. This observation is reminiscent of previous results in breast cancer and melanoma in which regulatory genes for the mesenchymal and de-differentiated phenotypes overlapped while those corresponding to epithelial and differentiated (melanocytic) phenotypes did not overlap and were tissue-specific (Klinke & Torang, 2020). Previous pan-cancer studies have also highlighted that downregulation of epithelial components and upregulation of mesenchymal features need not always be as strongly coupled as often assumed (Cook & Vanderhyden, 2020; Sahoo et al., 2022). Moreover, differences along these two axes need not be necessarily reflected at a transcriptional level (Norgard et al., 2021). Together, these observations highlight the need to analyze epithelial and mesenchymal axes independently, rather than as a conventional single metric for EMT.

Short duration of drug treatment can induce a NCSC phenotype that is highly mesenchymal (Fallahi-Sichani et al., 2017; Ramsdale et al., 2015). Previous studies in preclinical models have established that while the NCSC phenotype is attained by cells within 1-3 weeks of drug treatment, further de-differentiation to the undifferentiated phenotype occurs only after 8-12 weeks of drug treatment. Our study highlights the absence of a positive correlation between the mesenchymal signature and de-differentiation beyond the NCSC phenotype, suggesting that prolonged treatment (beyond the 3-week mark for the NCSC phenotype) induces further de-differentiation but no concomitant increase in mesenchymal status. This observation of the NCSC phenotype being the most mesenchymal is in accordance with melanocyte development. Neural crest cells are progenitors of melanocytes that undergo EMT during development to delaminate and migrate from the neural tube to the epidermis, where they lose their EMT signature and differentiate into melanocytes (Dupin & le Douarin, 2003; Tang et al., 2020; Wessely et al., 2021). Thus, the nonmonotonic variation in EMT during development (the initial increase during migration followed by decrease during differentiation) is recapitulated during treatment-induced de-differentiation. Therefore, we propose that the often-presumed overlap between the mesenchymal and invasive axes may arise from the lack of information for longer time scales (since most *in vitro* drug treatment studies are performed in under three weeks), and our assumptions about linearly increasing trends. However, increasing evidence suggests that maximum stemness is associated with hybrid E/M phenotypes rather than ‘extreme’ mesenchymal or epithelial phenotypes, suggesting that many such associations among axes of plasticity can be non-monotonic in nature (Grosse-Wilde et al., 2015; Kröger et al., 2019; Pasani et al., 2021).

Our results also indicate that metabolic programs can be linked either with the de-differentiation program or the mesenchymal program. OXPHOS and fatty acid oxidation are both indirectly regulated by MITF. In the case of OXPHOS, MITF regulates PGC-1α (Vazquez et al., 2013); in the fatty acid oxidation pathway, MITF regulates SCD (Vivas-García et al., 2020). MITF, which controls both metabolic pathways, decreases with increasing de-differentiation. This trend is explained by the decline in MITF associated with de-differentiation, in accordance with the MITF rheostat model (Rambow et al., 2019). On the other hand, glycolysis and HIF-1α signatures seem to be coregulated with the mesenchymal program. Previous studies in epithelial cancers have shown how well-established EMT transcription factors (EMT-TFs) regulate the metabolic profile of a cell and cause a switch to glycolysis (also called Warburg effect) (Youssef & Nieto, 2020). Interestingly, neural crest cells also display decay of glycolytic capabilities during differentiation into melanocytes (Zheng et al., 2016). Our analysis suggests that the metabolic state of a cell is closely linked to the transcriptional program governing it at a given time point. Thus, de-differentiation captures the transcriptional and metabolic states observed during melanocyte development.

Although our study focuses on melanoma, EMT-like phenotypic switching is also characteristic of other non-epithelial cancers and de-differentiation of melanocytes independent of malignant transformation. De-differentiation of melanocytes into pluripotent stem cells demonstrated a reduction in expression levels of E-Cadherin, an epithelial marker and similarities to mesenchymal stem cells (Vidács et al., 2022). Molecular subtypes of glioblastoma multiforme (GBM), a nonepithelial cancer, include the pro-neural, classical, and mesenchymal phenotypes, which exist along a spectrum of worsening prognosis (Fedele et al., 2019). Single-cell analysis reveals that these molecular subtypes recapitulate neurodevelopmental trajectories, with proneural cells forming a major composition of proliferative glial progenitor-like cells (Couturier et al., 2020; Phillips et al., 2006). A proneural-to-mesenchymal transition (PMT) is characterized by an increase in mesenchymal markers, such as SNAI1 and ZEB1. Interestingly, glioma stem cells (GSCs) exist as proneural GSCs and mesenchymal GSCs, which can give rise to the complete spectrum of intratumor heterogeneity, including the classical phenotype (Wang et al., 2019), reminiscent of epithelial and mesenchymal CSCs reported in breast cancer (Liu et al., 2013). Moreover, samples belonging to the classical subtype are depleted of pro-neural GSCs and enriched for mesenchymal GSCs, possibly suggesting that mesenchymal GSCs are more likely to give rise to the classical subtype. This trend strengthens the semi-independent nature of EMT and stemness as seen in epithelial cancers (Sahoo et al., 2022). Another study in GBM cell lines reports that loss of N-cadherin (a well-established mesenchymal marker) increases invasiveness (Camand et al., 2012), reinforcing the trends that increased migration and invasion is not an inexorable consequence of carcinoma-associated EMT (Schaeffer et al., 2014). These scenarios of non-overlapping behaviors in terms of invasiveness, stemness and EMT, seen both for epithelial and non-epithelial cancers, advocate for improving existing therapeutic strategies by targeting multiple axes of cellular plasticity simultaneously rather than individually.

Our study focuses on the overlap between the de-differentiation and the EMT axis during drug treatment in melanoma samples. However, de-differentiation is not the only trajectory taken up by cells during drug treatment. Cells can follow multiple paths to resistance, one of which is by attaining a hyper-pigmented phenotype (Goyal et al., 2021; Rambow et al., 2018; Su et al., 2020). The mapping of these trajectories and states to the E-M axis remains to be studied. In addition, another axis of cellular plasticity commonly associated with EMT is immune suppression and immune evasion. Previous studies have shown that the expression levels of programmed death-ligand 1 transmembrane protein (PD-L1) – a driver of immune evasion - does not increase monotonically with EMT (Sahoo et al., 2021). Consistently, in melanoma, the expected trend of worse response to anti-PD-1 therapy with increasing de-differentiation is not observed; rather, results from the CheckMate 038 clinical trial indicate that the NCSC phenotype is associated with a better outcome to immune checkpoint blockade therapy as compared to the melanocytic phenotype (Y. J. Kim et al., 2021). The extent of overlap between the axes of EMT, immune evasion, and de-differentiation require further study to design temporally-sequenced effective combination therapies that can shift the differentiation and EMT status of melanoma toward a less invasive and more immune activated state. Recent *in vitro* investigations in melanoma have shown proof-of-principle evidence of phenotypic plasticity driven drug resensitization as a mechanism underlying the beneficial impact of intermittent therapy (Kavran et al., 2022).

Overall, our transcriptomic data-based analysis highlights the partially overlapping nature of EMT with molecular phenotypes of de-differentiation and metabolism during drug treatment in melanoma. Better insights into these observed trends can be gained by mechanistic models to interrogate the coupling of the underlying regulatory networks that govern these processes. A better understanding of these dynamics can help identify more effective therapeutic strategies by fine-tuning the interval, sequence, and dosage of treatment (Goldman et al., 2015) and by developing combination and/or sequential therapeutic strategies. Our study provides a framework for studying these different axes of plasticity and heterogeneity independently and identifying the degree to which these axes overlap.

## Code Availability

All codes and scores generated for this paper can be found at: https://github.com/csbBSSE/EMT_Melanoma

## Acknowledgement

This work was supported by Ramanujan Fellowship awarded to MKJ by Science and Engineering Research Board (SERB), Department of Science and Technology (DST), Government of India (SB/S2/RJN-049/2018) and by Infosys Young Investigator award to MKJ supported by Infosys Foundation, Bangalore. MP is supported by KVPY fellowship (DST). JAS is supported by NIH 1R01CA233585-03, Department of Defense W81XWH-18-1-0189, and the Triangle Center for Evolutionary Medicine.

## Author contributions

MP and GR performed research, analyzed data and wrote the first draft of the manuscript. PT, NA, SM and AM performed research. JAS and MKJ conceived research and worked on manuscript revisions. MKJ supervised research.

## Supplementary Figures

**Fig S1.**
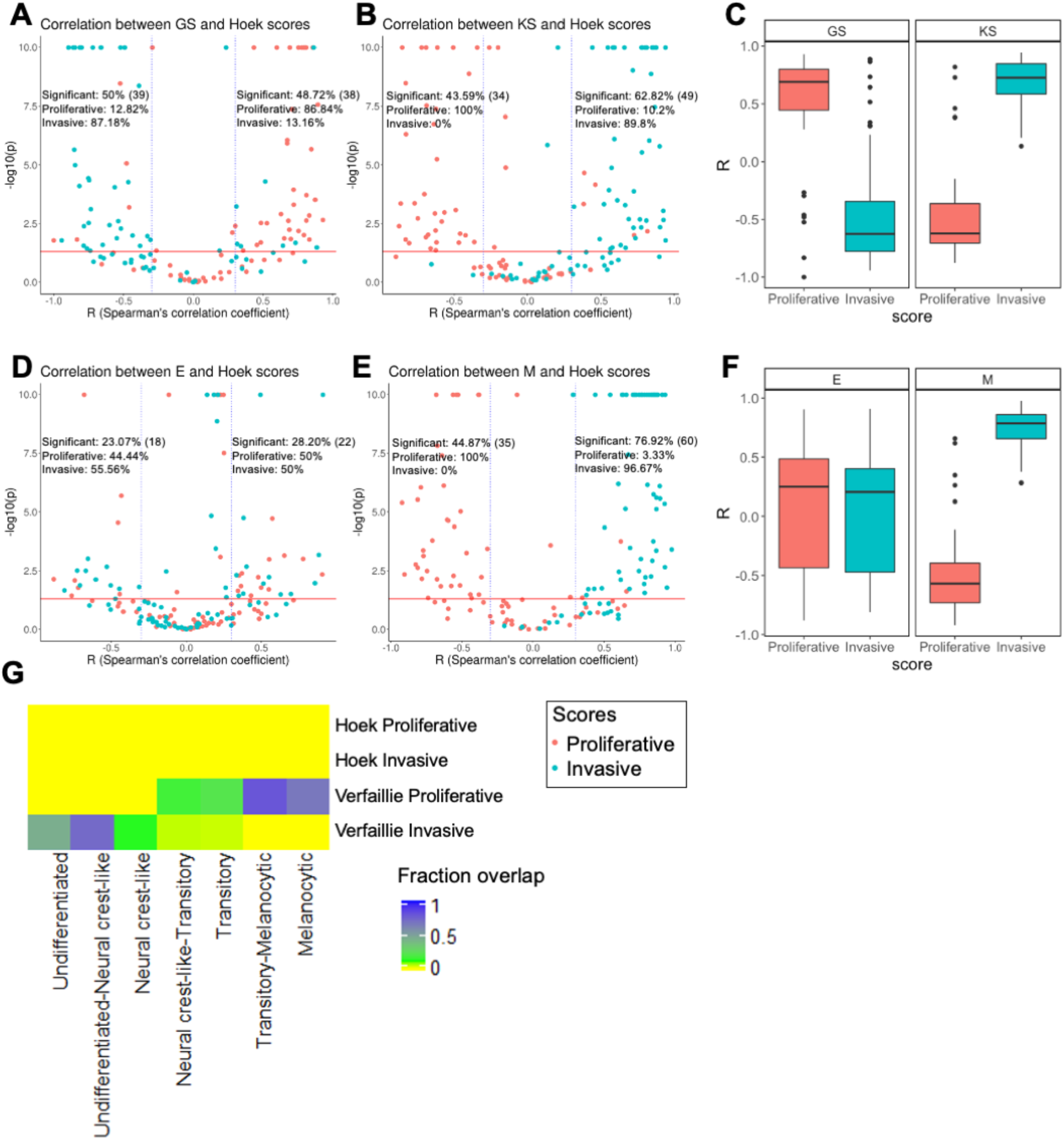
Volcano plots depicting the Spearman’s correlation coefficient and -log_10_(p-value) for Hoek proliferative and invasive gene set and **A**. 76GS EMT scoring metric **B**. KS EMT scoring metric **C**. Boxplot depicting range of correlation coefficients for KS and GS **D**. Epithelial gene set **E**. Mesenchymal gene set. **F**. Boxplot depicting range of correlation coefficients for E and M scores Inset labelled “Significant” is calculated as 100* number of significant points under specified cut-off /78. Absolute number of significant points for the specified cut of is mentioned in brackets. “Proliferative” and “Invasive” labels represent the percentage of significant correlations that are between the EMT score and proliferative score or invasive score, respectively. **G**. Extent of overlap between Verfaillie, Hoek gene sets (Proliferative and invasive phenotype gene sets) and Tsoi gene sets (intermediate phenotype gene sets)

**Fig S2.**
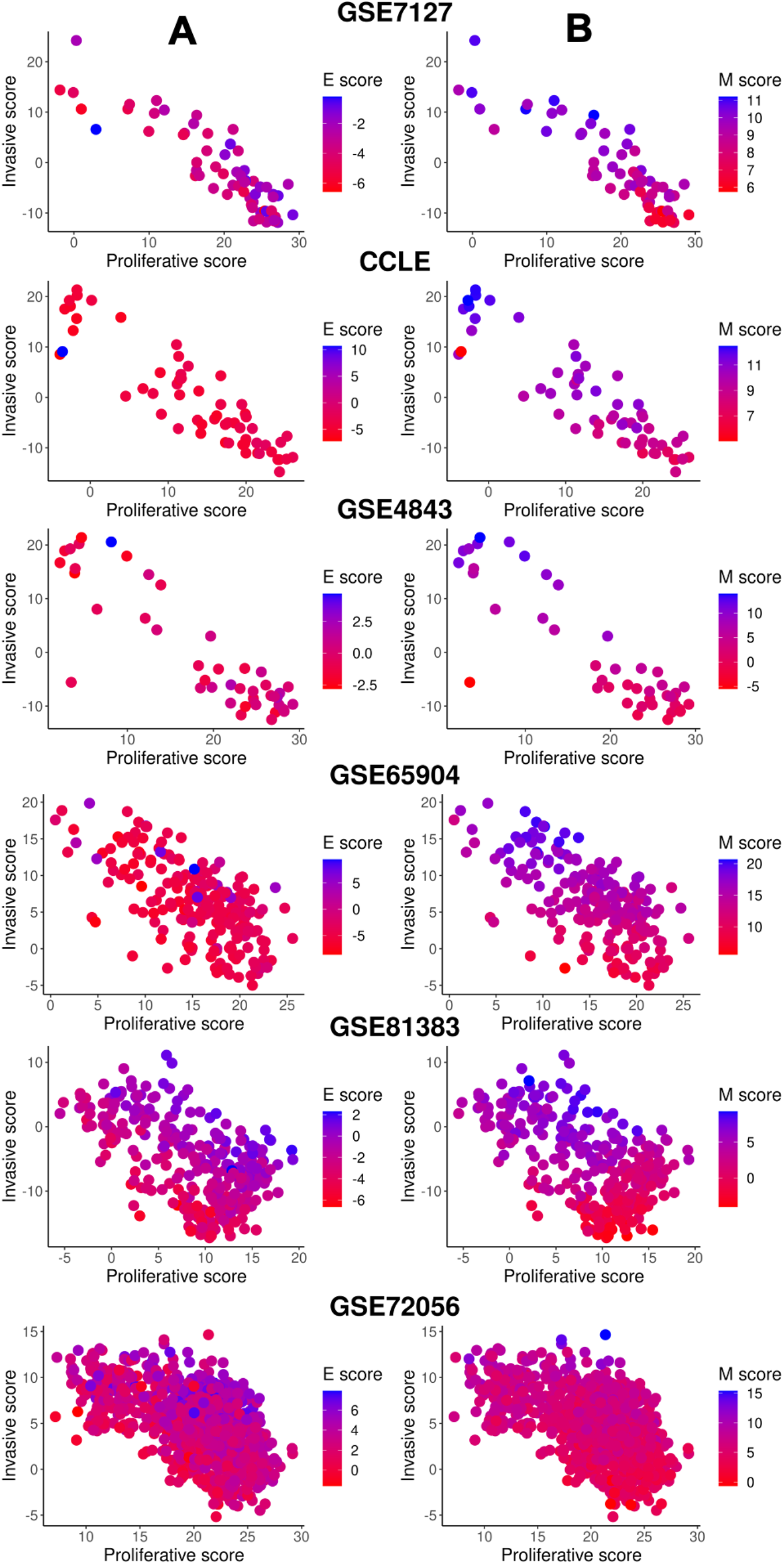
**A**. Two dimensional EMT plots depicting the variation of epithelial score gradient along the proliferative-invasive score axes. **B**. Two dimensional EMT plots depicting the variation of mesenchymal score gradient along the proliferative-invasive score axes.

**Fig S3.**
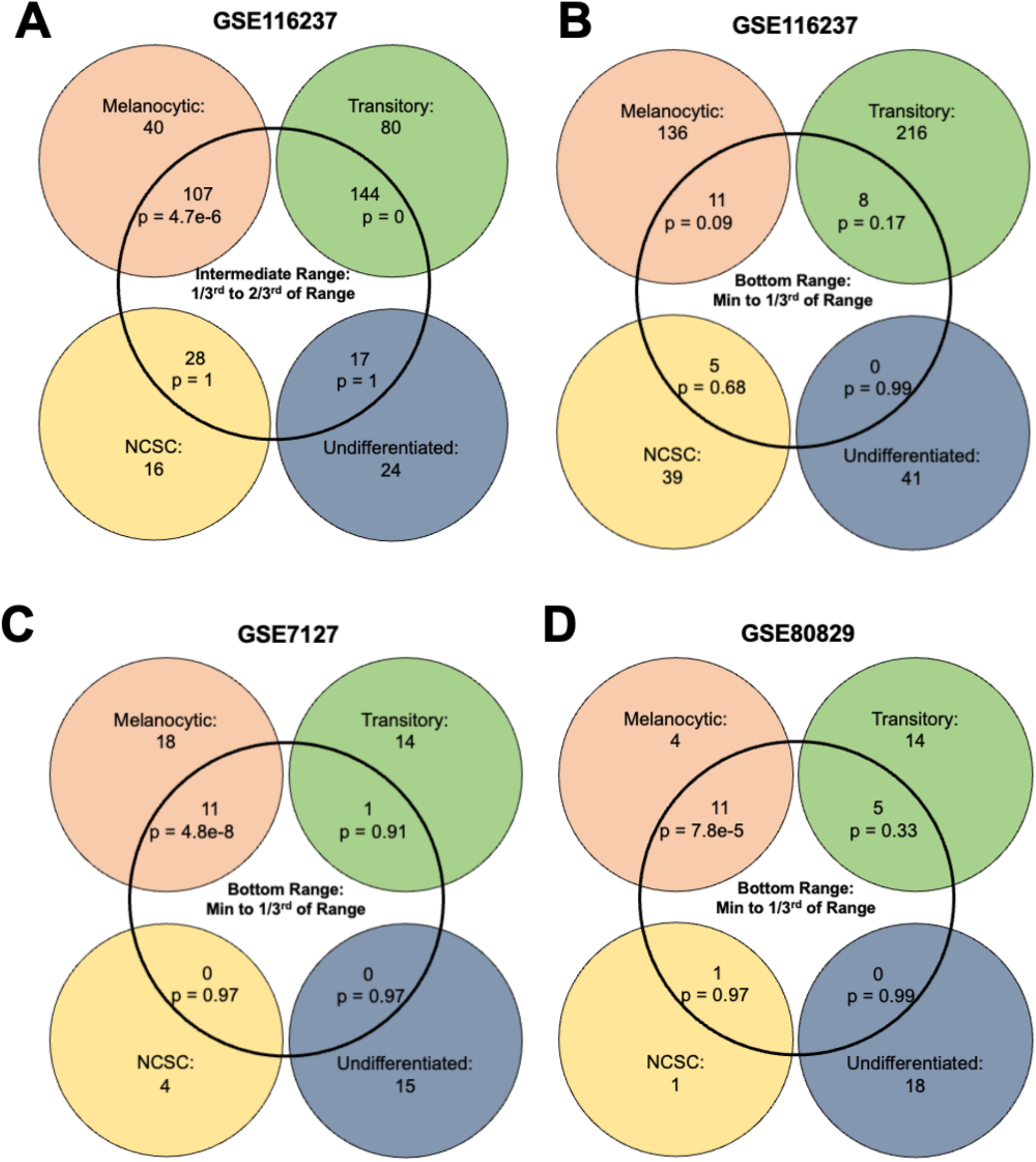
Venn diagram depicting **A**. The intersection of the four phenotype scores of samples and intermediate M scores in GSE116237. p represents p-value for the conditional probability that a sample belongs to the phenotype given that they lie in the intermediate M score range. The intersection of the four phenotype scores of samples and bottom M scores in **B**. GSE116237 **C**. GSE7127 **D**.GSE80829. p represents p-value for the conditional probability that a sample belongs to the phenotype given that they lie in the bottom M score range.

**Fig S4.**
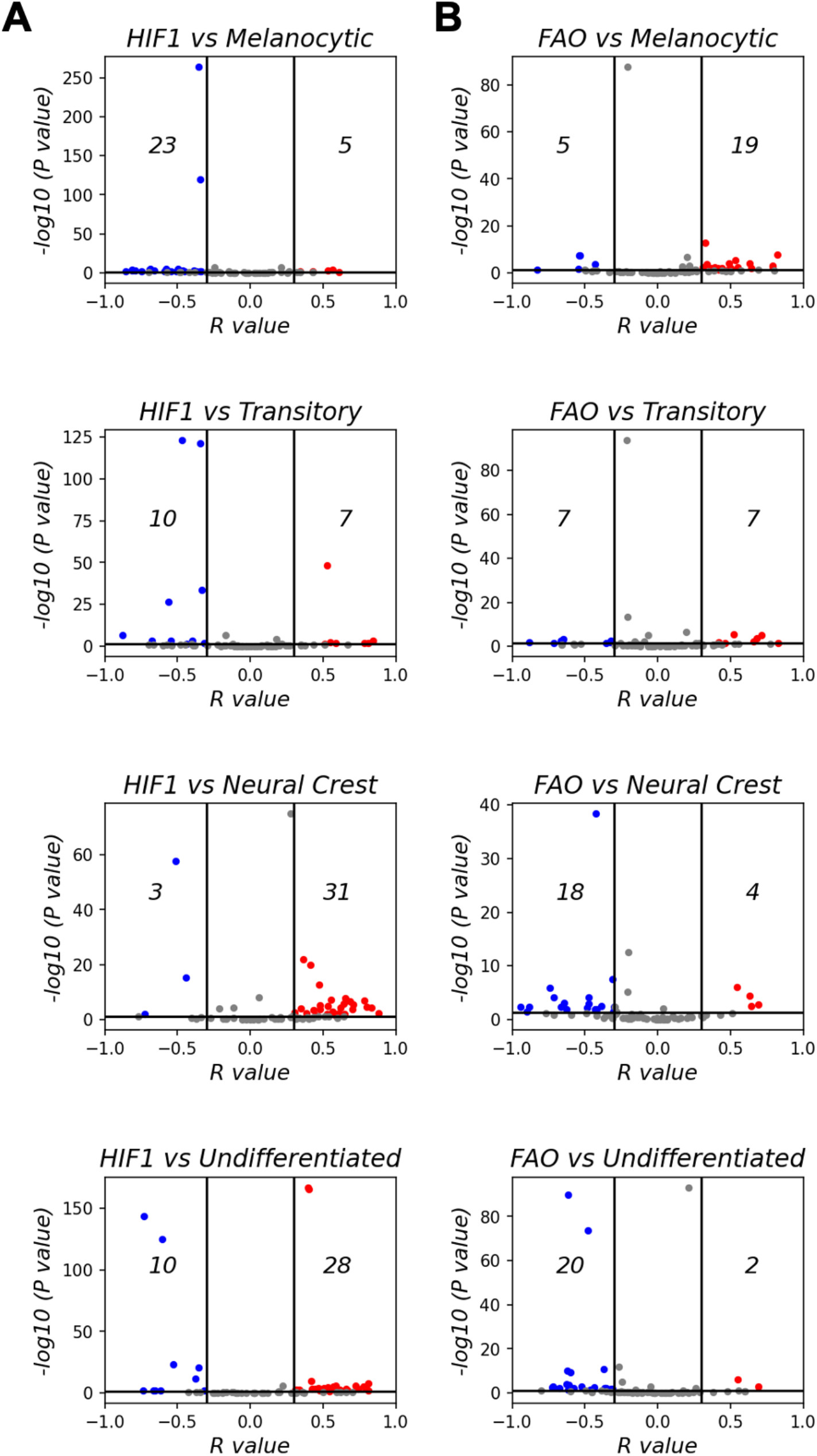
Volcano plots depicting the Spearman’s correlation coefficient and -log_10_(p-value) for **A**. HIF1 signature and Tsoi gene set. **B**. Fatty acid oxidation (FAO) gene set and Tsoi gene set.

